# Genome size and nucleotide skews as predictors of bacterial growth rate

**DOI:** 10.1101/2025.09.17.676822

**Authors:** Parthasarathi Sahu, Sashikanta Barik, Koushik Ghosh, Hemachander Subramanian

**Affiliations:** Department of Physics, National Institute of Technology Durgapur, India - 713209

**Keywords:** Replication time, Genome evolution, Replichore organization, Fork speed, Nucleotide skew, Asymmetric Cooperativity Model

## Abstract

Bacterial growth rates are constrained by genome replication, yet the role of replication kinetics in bacterial growth rates remains incompletely understood. Here, we examine if genome size, replichore organization, and nucleotide compositional asymmetry are reasonable predictors of bacterial doubling times. In free-living bacteria, both genome size and the length of the longest replichore are found to correlate positively with doubling time, pointing to an influence of replication dynamics on bacterial growth rates. Moreover, fast-growing bacteria are shown to exhibit stronger nucleotide compositional skew. Incorporating skew into the model substantially improves predictive accuracy, suggesting that compositional asymmetry in genomes may facilitate replication fork progression and thereby enhance growth rates. Based on these observations, we speculate that nucleotide skew may play a potential adaptive role in bacterial genome replication. To assess whether the observed association between genome architecture and growth rate reflects an evolutionary signature or a mechanistic link, we reconstructed ancestral states and found that the model fits ancestral traits more strongly, with predictive strength (*R*^2^) decreasing progressively along the evolutionary tree as successive speciations occur. We speculate that this association has been stronger early in bacterial evolution and became subsequently screened as organisms diversified and increased in ecological and physiological complexity.

## 1 Introduction

The persistence of *compact* genomes in bacteria, despite billions of years of evolution, has been explained by genome streamlining, where natural selection favors the reduction of genome size [1, 2, 3, 4]. For such a reduction to be maintained, it must confer measurable fitness advantages to the organism. Several mechanisms have been proposed: First, smaller genomes require fewer resources for maintenance and expression. Thus, in nutrient-limited environments and in species with large effective population sizes, selection is expected to favor reduced genomes [5, 6, 7]. Second, a reduction in genome size often correlates with smaller cell volume, which increases the surface-to-volume ratio and thereby enhances metabolic efficiency [7, 8]. Third, shorter genomes reduce the time required for replication, which could potentially accelerate cell division and improve competitive fitness. While the first two explanations, nutrient economy and enhanced metabolic efficiency, are widely cited to support streamlining theory, the third explanation has been more controversial. Earlier studies reported little to no correlation between bacterial generation time and genome size, leading to the dismissal of replication time as a significant driver of genome reduction [9, 10, 11, 12].

In this work, we revisit this question. We analyze two sets of bacterial genomes (*n* = 367 and *n* = 180) [11, 13] and their replichore organization to test the influence of genome architecture on the organism’s growth rate.

## Methods

This study examines how bacterial genome architecture influences replication dynamics and, consequently, organismal growth rates. We correlate growth rates with genome size, replichore size, and nucleotide compositional skew through linear regression models. Two datasets of bacterial growth rates are used: (i) doubling time normalized across reported data, Dataset-I [13], and (ii) minimal doubling time under rapid growth conditions, Dataset-II [11]. Corresponding genomic sequences are retrieved from the NCBI Genome Database, and their accession numbers are provided in Supplementary Data. Only species with both doubling time and genome sequence data available, i.e., 367 in Dataset-I and 180 in Dataset-II, are included in the analysis.

Throughout this study, statistical significance is assessed at *p* < 0.05, and effect sizes are interpreted using the coefficient of determination (*R*^2^), with *R*^2^ < 0.1 considered negligible, 0.1 ≤ *R*^2^ < 0.2 weak, and 0.2 ≤ *R*^2^ < 0.3 moderate.

### Identification of replication origin and terminus

Nucleotide compositional asymmetries have long been used to infer replication origins and termini [14, 15, 16, 17]. In this study, we employ the purine–pyrimidine skew (RY-skew), which quantifies imbalances between purines (*R* = {*A, G*}) and pyrimidines (*Y* = {*C, T*}) across the genome. The RY-cumulative skew is calculated as:

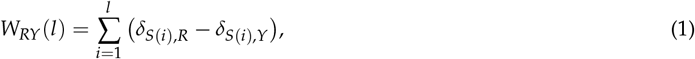

where *S* represents a genomic sequence of length *G, l* is the genomic position index running from 1 to *G* and *δ* is the Kronecker delta.

To find the origins and termini of any given sequence, we find the genomic segment that exhibits maximum RY-disparity (Fig. 1). Out of the two extremes of this segment, one falls in a valley position in the cumulative skew diagram, designated as the ‘replication origin’, and the other falls on a peak and is designated as the ‘terminus’, thereby partitioning the chromosome into two replichores [14]. A representative cumulative skew profile is shown in Fig. 1. All analyses were implemented in MATLAB, and custom scripts are publicly available (see Supplementary Information).

**Figure 1.**
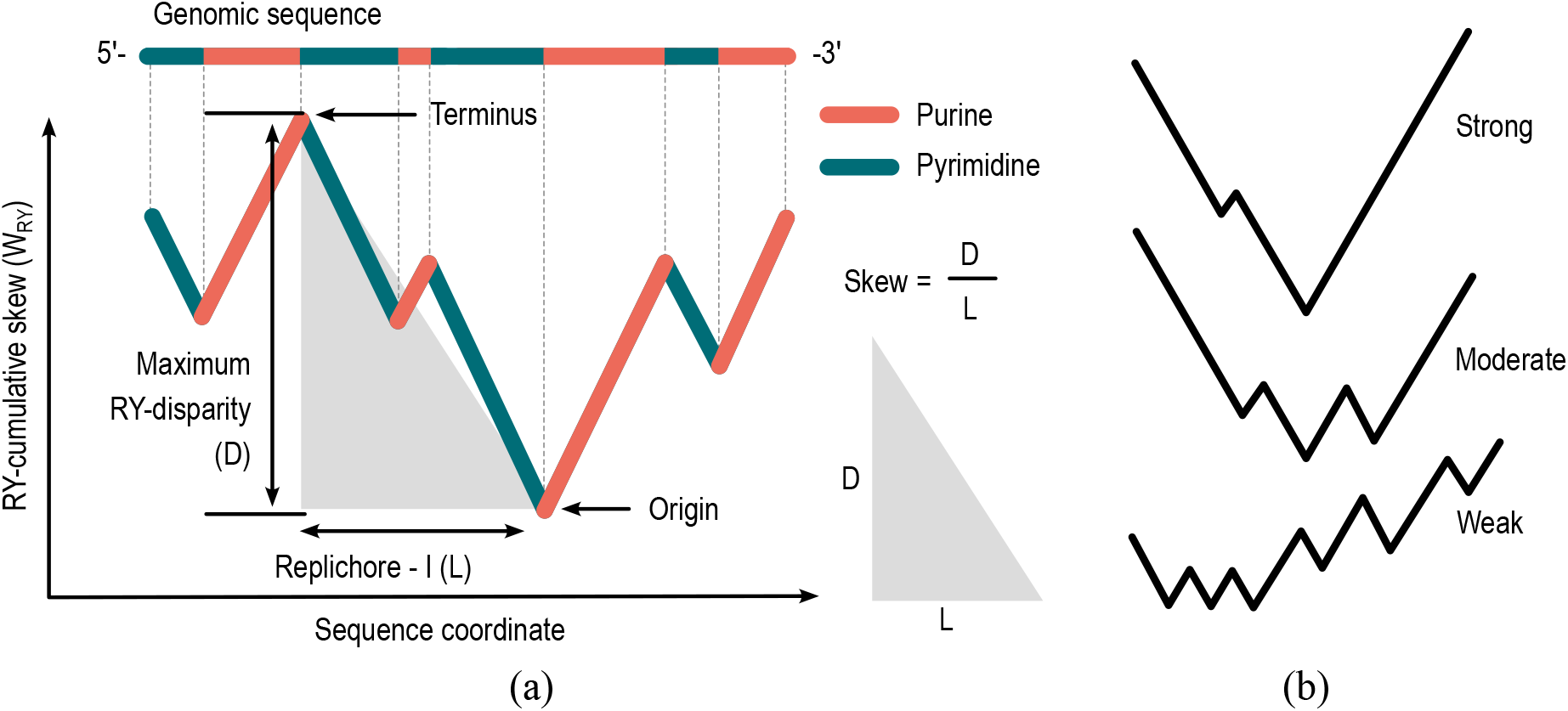
Identification of replication origin and terminus. (a) Cumulative RY-skew (*W*_*RY*_) for a representative genome segment, calculated using Eq. 1. The positions corresponding to the minimum and maximum cumulative RY-skews were used to predict the replication origin and terminus, respectively, thereby dividing the genome into two replichores. The ratio of maximum nucleotide disparity, D, and the corresponding replichore length, *L*, is defined as “skew”, a metric that measures the density of RY asymmetry in a replichore. (b) Cumulative skew profiles for three representative genomes, illustrating variation in skew strength categorized as “strong”, “moderate”, and “weak”.

## Results

We investigate whether the size and sequence structure of the genome can predict bacterial doubling time. To this end, we exclusively used genomic sequence characteristics, such as the size and organization of chromosomes and replichores, and their nucleotide compositional skew, to estimate the replication time of the genome and examined whether these characteristics correlate with bacterial doubling time using four regression models.

### Genome size and growth rate

As a first approximation, we modeled replication time as directly proportional to genome size, assuming a constant replication rate without parallelization. Here, by genome size, we imply all chromosomes present within a single bacterial cell, primary, and secondary chromosomes, if any. In this simplified form,

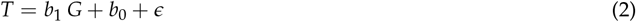

where *T* is replication time, *G* is genome size, and *b*_0_ and *b*_1_ are the regression coefficients of the model.

In agreement with previous reports [9, 10, 11, 12], we found a negligible or an absence of correlation between genome size and doubling time when considering the full dataset, *R*^2^ = − 0.00 for Dataset-I and *R*^2^ = −0.04 for Dataset-II, see Fig. 2, Table 1 Model-I. Excluding obligate intracellular species, which are strictly host-dependent bacteria and hence largely insulated from nutrient competition [18], we found an increase in correlation, although it remained insignificant (*p* > 0.05) in Dataset-II. However, when restricting the analysis to free-living bacteria, further excluding facultative host-dependent bacteria, the correlation became more pronounced, *R*^2^ = 0.09 in Dataset-I, suggesting that in *competitive* environments, bacterial doubling time is influenced by genome size. This weak correlation arises from a simplified model that ignores parallel replication across chromosomes and replichores, thus overestimates replication time relative to biological reality [19, 20]. On excluding secondary chromosomes and correlating only the size of the *primary* chromosome to the doubling time, we observe a better correlation than that with the total genome size, with *R*^2^ = 0.01 for the entire dataset, and *R*^2^ = 0.11 for free-living bacteria in Dataset-I and *R*^2^ = − 0.03 (entire dataset) and *R*^2^ = 0.06 (free-living) in Dataset-II, see Fig. 3, Table 1, Model-II.

**Table 1.** The bacterial doubling time is correlated with 4 different predictors. The *R*^2^ and *p*-values for two datasets under different replication models are presented.

| Dataset | Subset | Model I |  | Model II |  | Model III |  | Model IV |  |
| --- | --- | --- | --- | --- | --- | --- | --- | --- | --- |
| | | No parallelization<br>($T \propto$ genome size) | | Parallelization across<br>chromosomes<br>($T \propto$ chromosome<br>size) | | Parallelization across<br>chromosomes &<br>replichores<br>($T \propto$ longest<br>replichore) | | Fork speed depends<br>on RY-skew<br>($T \propto$ replicore<br>length/skew) | |
| | | $R^2$ | $p$ | $R^2$ | $p$ | $R^2$ | $p$ | $R^2$ | $p$ |
| Dataset I<br>Total 367<br>(Madin et<br>al. [13]) | All organisms ($n = 367$) | 0.00 | <u>3e-01</u> | 0.01 | <u>8e-02</u> | 0.03 | 7e-04 | 0.15 | 3e-14 |
| | Excluding Obligate<br>Intracellular<br>($n = 346$) | 0.02 | 4e-03 | 0.04 | 3e-04 | 0.09 | 3e-08 | 0.19 | 5e-17 |
| | Free-living ($n = 187$) | 0.09 | 2e-05 | 0.11 | 4e-06 | 0.15 | 4e-08 | 0.18 | 1e-09 |
| Dataset II<br>Total 180<br>(Vieira et<br>al. [11]) | All organisms ($n = 180$) | -0.04 | 7e-03 | -0.03 | 3e-02 | -0.01 | 2e-01 | 0.04 | 6e-03 |
| | Excluding Obligate<br>Intracellular<br>($n = 159$) | 0.01 | <u>3e-01</u> | 0.02 | <u>9e-02</u> | 0.05 | 7e-03 | 0.20 | 2e-09 |
| | Free-living ($n = 104$) | 0.03 | <u>7e-02</u> | 0.06 | 1e-02 | 0.11 | 5e-04 | 0.30 | 2e-09 |

**Figure 2.**
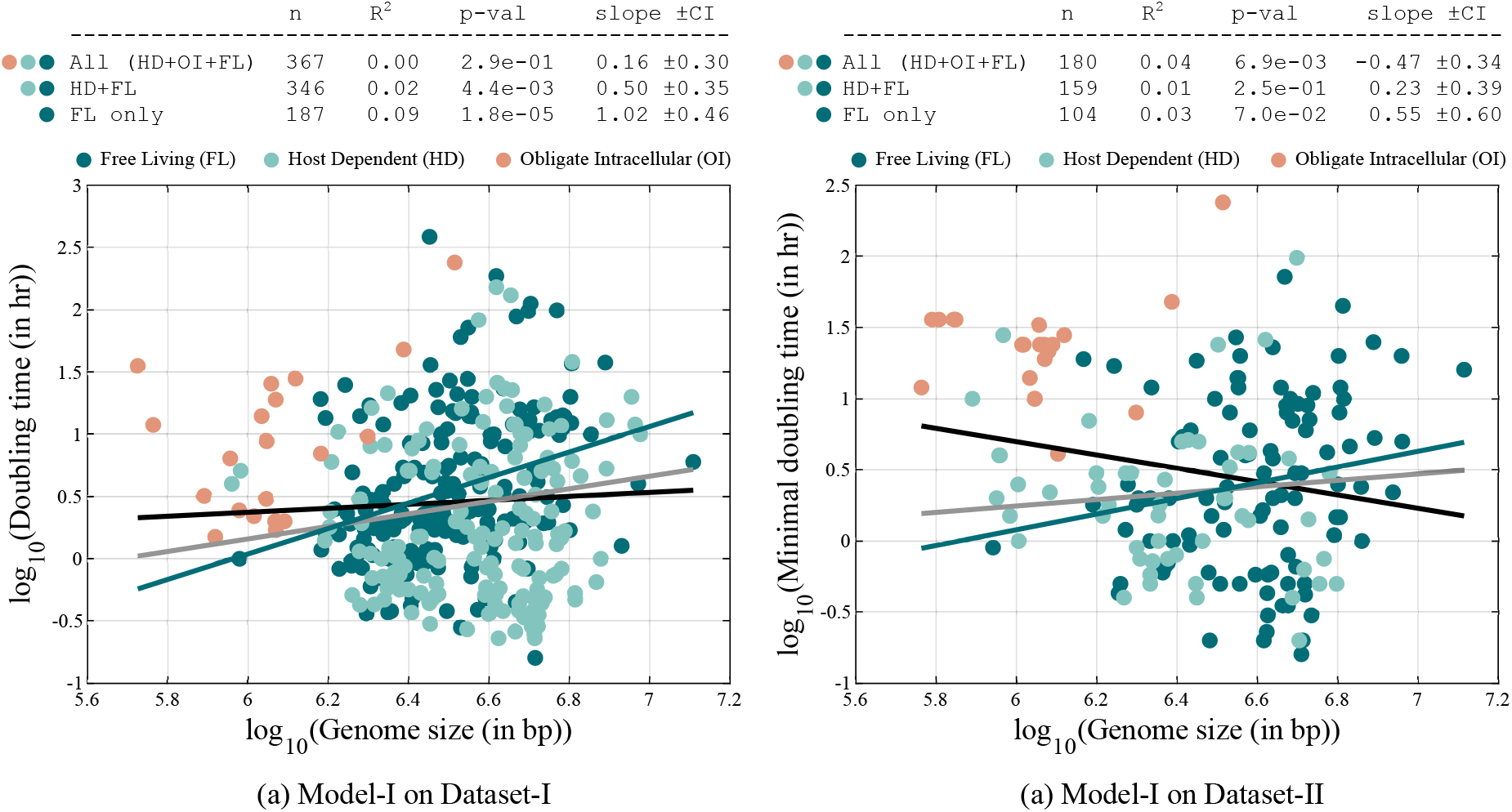
Relationship between bacterial doubling time and genome size in (a) Dataset-I and (b) Dataset-II. An absence of correlation in Dataset-I and a negligible negative correlation is observed in the full data set. Restricting the analysis to only free-living bacteria reveals a negligible positive correlation in Dataset-I. Regression lines are shown in black for the full dataset, gray for the dataset excluding obligate intracellular species, and green for free-living bacteria only. The same color scheme is followed throughout all figures in this manuscript.

**Figure 3.**
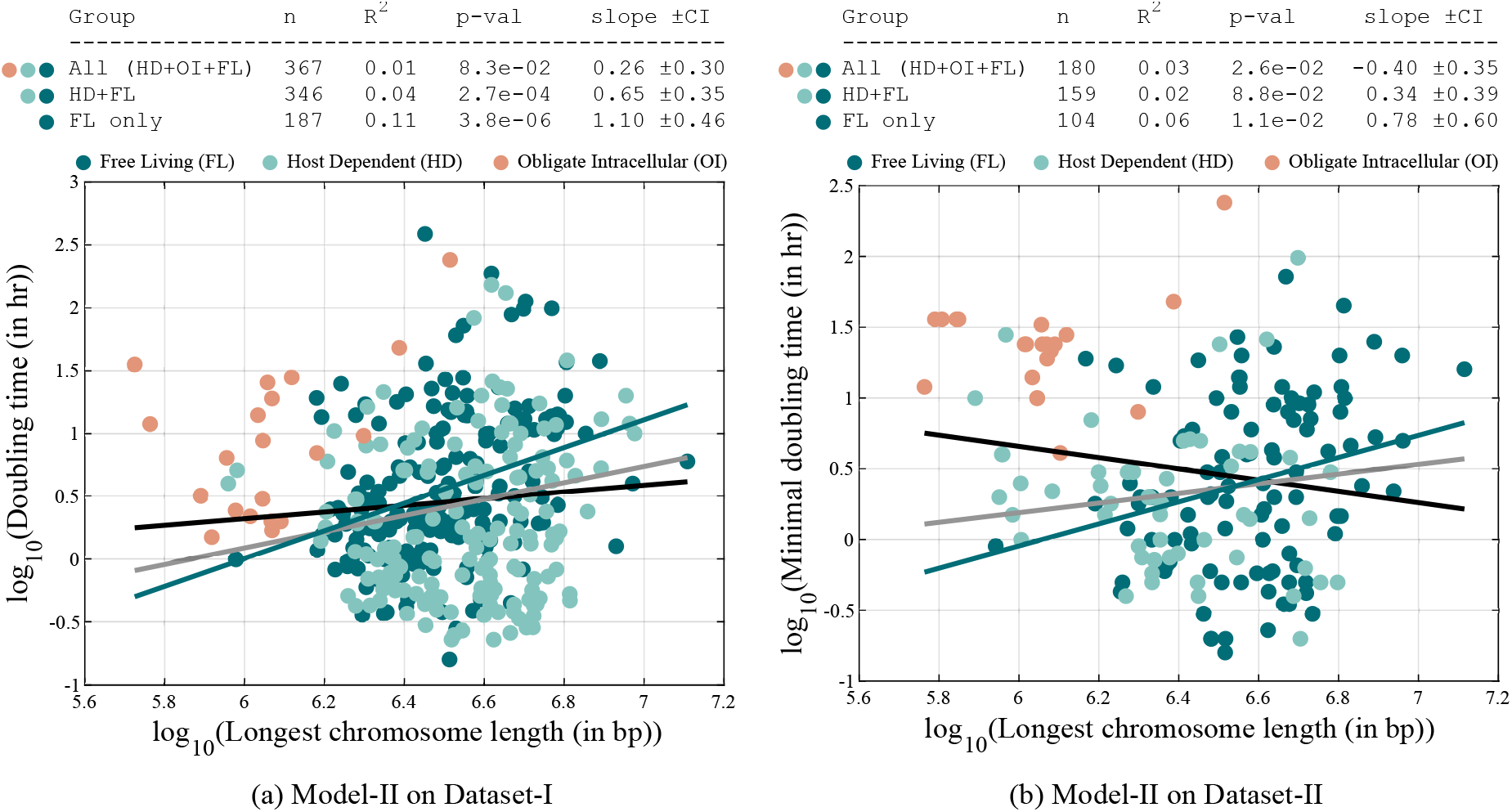
Relationship between bacterial doubling time and chromosome size in (a) Dataset-I and in (b) Dataset-II. The correlation between doubling time and chromosome size improves compared to that with total genome size. As the secondary chromosomes, ploidies are replicated in parallel [20]; the longest chromosome, typically the primary one, dictates the replication time.

### Length of the longest replichore and growth rate

Following the previous analysis, assuming parallel replication across chromosomes and also the replichores, the effective replication time is determined by the longest replichore. Accordingly, we refined the model to

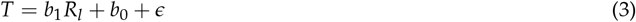

where *R*_*l*_ is the length of the longest replichore, identified by using purine-pyrimidine compositional skew in the primary chromosome.

Incorporating replication parallelization improved the model’s prediction of doubling time with *R*^2^ = 0.03 for all organisms and *R*^2^ = 0.15 for free-living bacteria in Dataset-I and *R*^2^ = − 0.01 for the entire dataset and *R*^2^ = 0.11 for free-living bacteria in Dataset-II, see Fig. 4, Table 1, Model-III.

**Figure 4.**
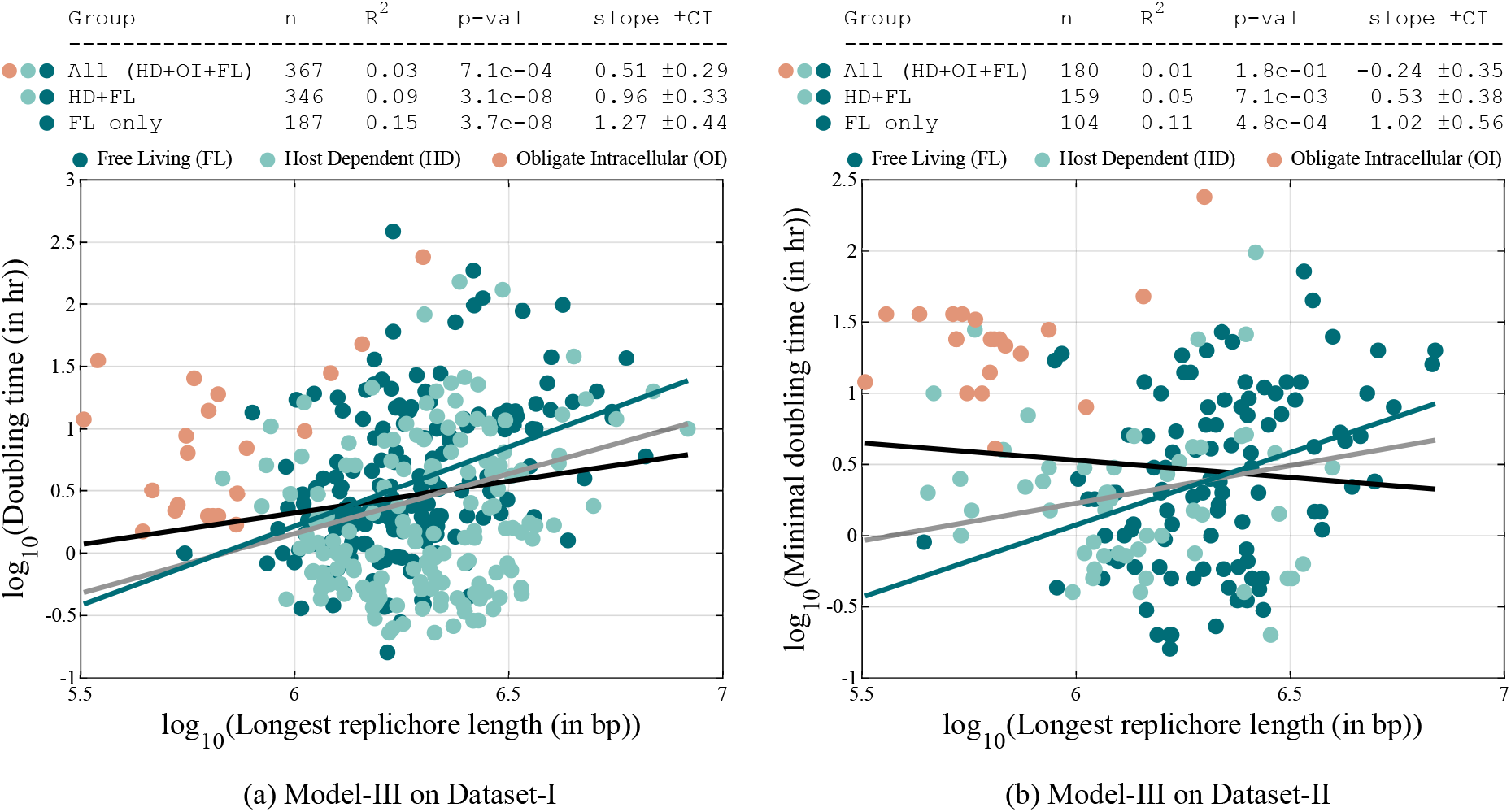
Relationship between bacterial doubling time and the length of the longest replichore in (a) Dataset-I and in (b) Dataset-II. Accounting for replication parallelization leads to a better correlation than when using total genome size, highlighting the importance of replichore architecture in shaping replication dynamics.

### Effect of skew on replication kinetics

Inspired by our previous DNA unzipping simulation [21], which suggested that homogeneous sequences (compositionally skewed sequences) facilitate rapid sequential strand separation, we examined whether such nucleotide compositional asymmetry in bacteria influences their replication dynamics. We define skew as the slope of the genomic segment exhibiting maximum RY-disparity (see Fig. 1). Aligned with previous reports [15, 22], we observed that fast-growing bacteria tend to exhibit stronger purine–pyrimidine skew in their sequence (Fig. 5).

**Figure 5.**
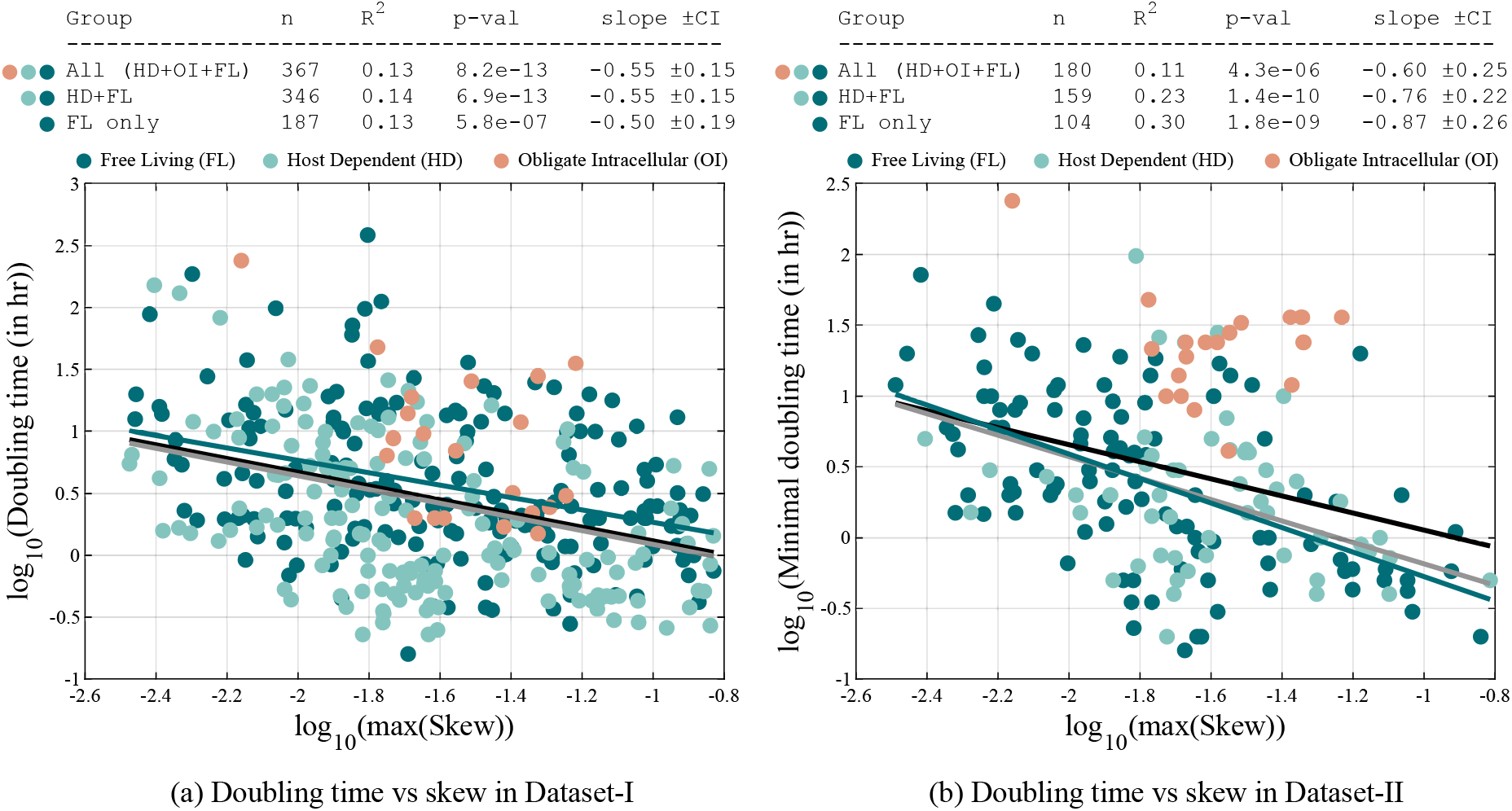
Relationship between bacterial doubling time and maximum purine–pyrimidine skew of the genome in (a) Dataset-I and in (b) Dataset-II. Faster-growing bacteria exhibit stronger compositional skew, suggesting a potential role of skew in replication kinetics.

Based on this observation, we hypothesize that nucleotide skew may enhance replication fork kinetics and, consequently, affect bacterial growth rates. To investigate this, we collected reported replication fork speed data for 10 distinct species from 25 different literature sources (see Supplementary Information), and their correlation with nucleotide skew is shown in Fig. 6(b). Due to the limited data size, the co-variation of replication fork speed with nucleotide skew remains inconclusive and requires further analysis.

**Figure 6.**
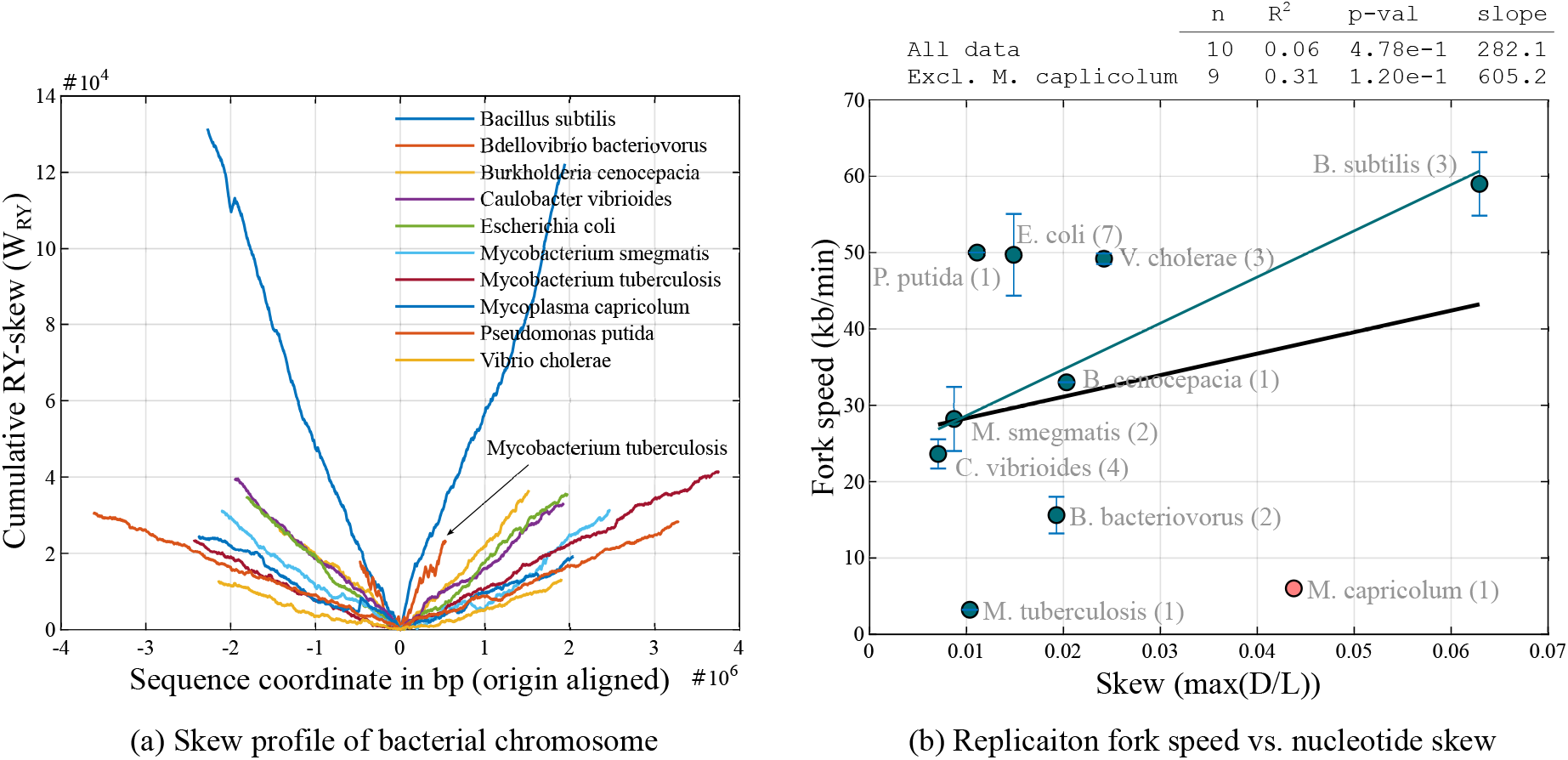
Correlation between reported replication-fork speed and the maximum purine–pyrimidine skew of the replichores. The principal outlier, *Mycoplasma capricolum*, has a highly reduced genome with very low GC content (¡25%), has lost numerous replication-associated proteins, and lacks *de novo* nucleotide synthesis, relying instead on nucleotide interconversion [23, 24]. Although a moderate correlation is observed when this outlier is excluded, the overall relationship remains inconclusive due to the limited sample size.

To model the above consideration, we introduce the assumption that the replication fork speed scales with nucleotide skew. The predictor of replication time is now defined as the maximum ratio of replichore length to its corresponding skew (representing fork speed), calculated across both replichores, Model-IV. Under this assumption, replication time can be approximated as

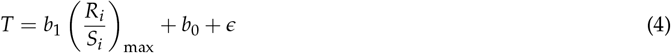

where (*R*_*i*_/*S*_*i*_)_max_ is the maximum of the length-to-skew ratios across the two replichores. Incorporating skew into the regression model improved predictive performance compared with the previous model using replichore size alone, yielding *R*^2^ = 0.15 for the total data set and *R*^2^ = 0.18 for free-living bacteria in Dataset-I and *R*^2^ = 0.04 for the entire dataset and *R*^2^ = 0.30 for free-living bacteria in Dataset-II, see Fig. 7 and Table 1, Model-IV.

**Figure 7.**
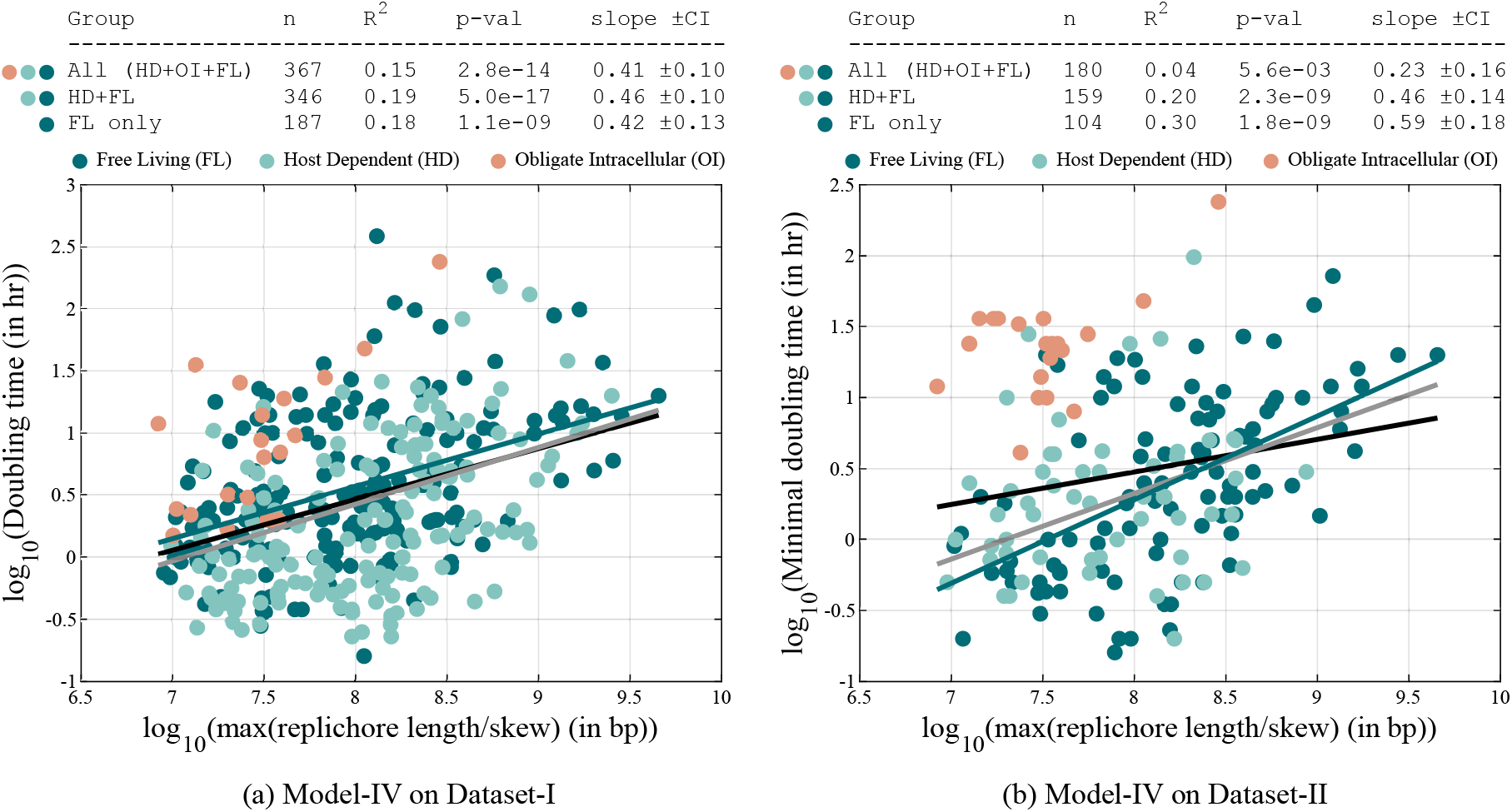
Predicted versus observed doubling time under the skew-adjusted model in (a) Dataset-I and in (b) Dataset-II, where replication time is modeled as the ratio of replichore length and the fork speed, with the latter assumed to be proportional to skew. Including nucleotide skew strengthens the predictive power compared to models based solely on genome size or replichore length.

### Test for phylogeny and other confounding factors

We performed additional analyses on Model-IV of Dataset-I to assess the potential influence of confounding factors on the observed correlation. We examined genomic GC content, optimum pH, growth temperature, and optimal temperature using the data provided [13] and found that none of these variables acted as confounders; see the Supplementary Information for the partial correlation analysis. We further observed that although within-clade correlations between the two traits are mostly absent or moderate, correlations between clade-level mean traits, i.e., the geometric mean of the trait values among the extant species within each clade, are noticeably stronger; see Supplementary Information.

To assess the influence of phylogeny on the observed correlations, we assembled divergence-time estimates from TimeTree [25] for 323 of the 367 species in Dataset-I; see Supplementary Information for the phylogenetic tree and Supplementary Data for the Newick file. Two terminals with zero branch lengths were removed to avoid singularities in the phylogenetic covariance matrix *C*. In addition, we applied a Pagel’s *λ* transformation [26] to the branch lengths to reduce the ill-conditioning of *C* (initial condition number ∼ 4 × 10^10^). Incorporating phylogenetic structure via Phylogenetic Generalized Least Squares (PGLS) [27] eliminated the correlation (PGLS *R*^2^ ∼ 0.05) detected under Ordinary Least Squares (OLS) in Model-IV (Table 2).

**Table 2.**
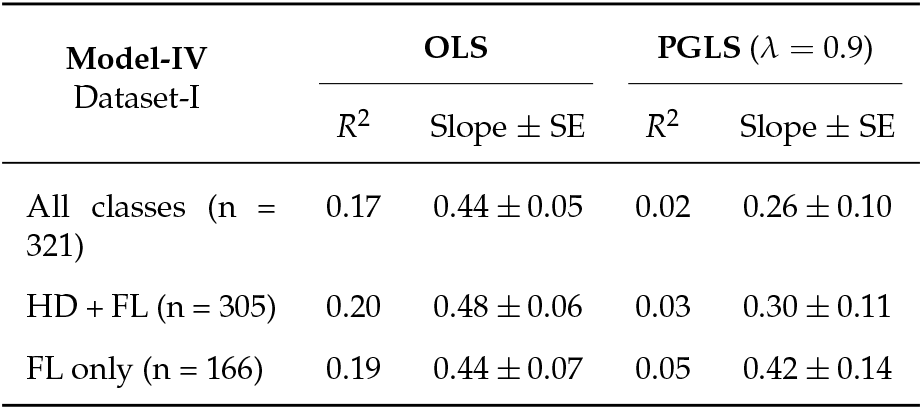
Comparison of ordinary least squares (OLS) and phylogenetic generalized least squares (PGLS) regression statistics on Dataset-I for the relationship between skew normalized replichore length (predictor) and doubling time (response).

To better understand this outcome, we quantified phylogenetic signal in both the predictor and the response using Blomberg’s *K* [28], after rescaling the covariance matrix *C*. We found that the predictor, i.e., skew-normalized replichore length, exhibits a strong phylogenetic signal (*K* ≈ 1.5), implying that close relatives are more similar than expected under Brownian motion evolution, whereas the response, doubling time, and its residuals show a weak signal (*K* ≈ 0.49). Such data are reported to reduce the reliability and statistical power of PGLS analyses [29, 30]. We further examined the influence of phylogeny on Model-IV by estimating the model’s performance in early evolutionary time periods.

To examine ancestral patterns, we estimate trait values for all ancestral lineages (320 internal nodes) in the Dataset-I phylogeny using the observed traits of the 321 extant species. The ancestral states correspond to the set of internal node values that maximize the likelihood of the observed tip data under a Brownian-motion model of trait evolution. These maximum-likelihood (ML) ancestral states can be computed using a closed-form solution in which each internal node value is expressed as a weighted average of the descendant tip values, with weights determined by the variances implied by the branch lengths [31, 32]. We implemented this algorithm to obtain the ML estimates of trait values at all ancestral nodes; the MATLAB script is provided in the Supplementary Data.

The reconstructed ancestral data of the predictor and response at each time slice is then fitted to Model-IV, and the resulting *R*^2^ and *p*-value are shown as functions of evolutionary time in Fig. 8(b). A consistent increase in *R*^2^ is observed as the evolutionary time deepens, indicating that the association between genome architecture and doubling time in bacteria was stronger on the primitive Earth. The weaker correlations observed in present-day species are likely the result of accumulated lineage-specific divergence and adaptation that progressively obscure the historical signal. The scatter plots of the model fit across 50 time slices are compiled into a video (see Supplementary Information), providing a more intuitive visualization of how the association evolves over deep time.

**Figure 8.**
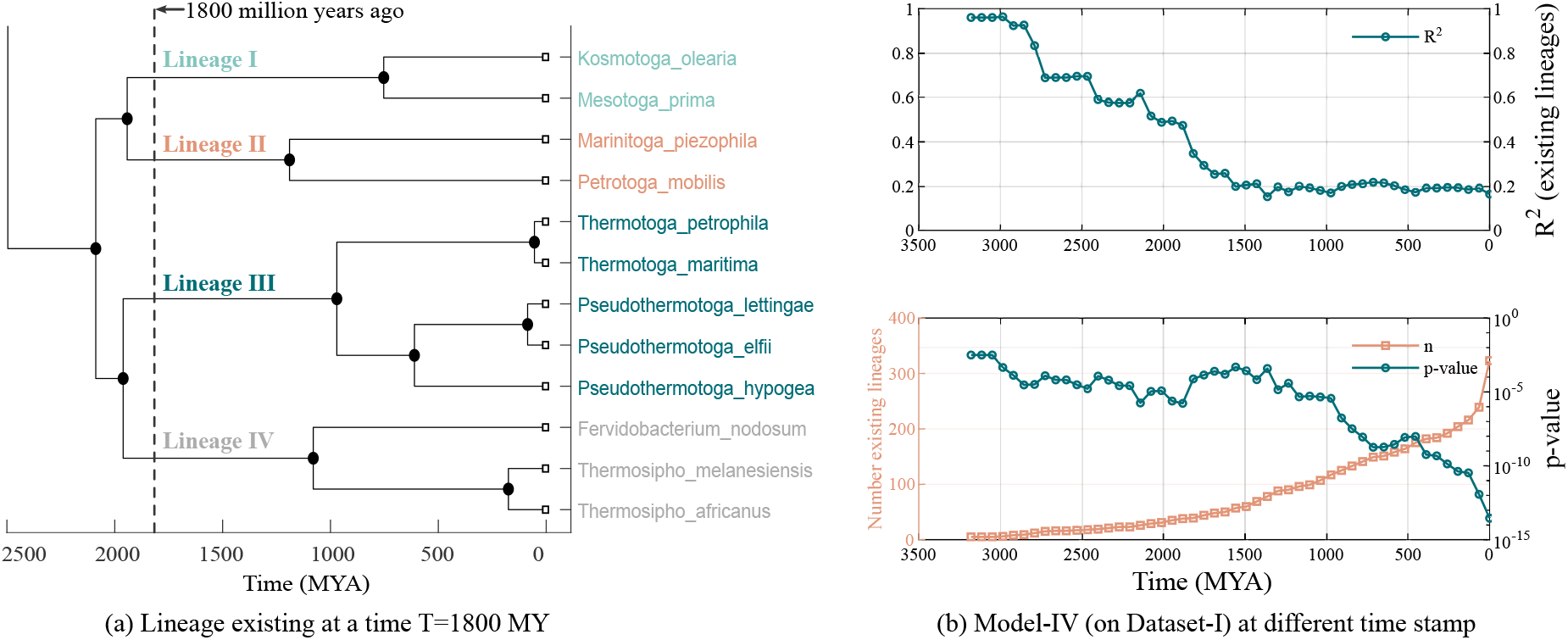
Performance of the model over evolutionary time. (a) The ancestral lineages existing at a given evolutionary time slice (shown here at 1800 million years ago, MYA). The predictor and response data for these ancestors are estimated under a Brownian motion model of evolution. (b) Performance of Model-IV is evaluated across evolutionary time, from the present day back to 3200 MYA. For each time slice, the reconstructed ancestral dataset is fitted to the regression model, and the resulting *R*^2^ and *p*-value of the model are plotted. Going back in time, the decrease in model significance, as indicated by the high *p*-value, is a result of a decrease in sample size, i.e., the number of clades. The progressive increase in *R*^2^ with deeper evolutionary time suggests that the underlying correlation between traits was stronger in the past and has been progressively obscured by lineage-specific divergence and accumulated stochastic noise over evolutionary history.

## Discussion

This study examines the relationship between bacterial genome architecture — specifically, genome size, replichore length, and nucleotide compositional skew — and bacterial growth rate. By estimating replication times from these genomic features and comparing them with empirically reported doubling times across a broad set of bacterial species, we present the following three principal outcomes.

First, genome size alone exhibits only a negligible association with doubling time, and this association differs markedly between ecological categories, i.e., obligate intracellular and free-living bacteria show distinct regression slopes and intercepts. Among organisms of comparable genome size, obligate intracellular bacteria generally grow more slowly, likely reflecting the reduced competitive pressures characteristic of their stable host-associated niches. We also observe that the association of genome size is not limited to bacteria; our analysis of archaeal genomes (*n* = 153) also revealed a weak positive correlation (*R*^2^ = 0.12, *p* < 1*e* − 05) between genome size and their doubling time (see Supplementary Information). Second, the length of the longest replichore is a better predictor of growth rate than total genome size, because DNA replication proceeds bidirectionally from the origin, and the slowest-replicating replichore constrains the overall replication time. Third, incorporating nucleotide compositional skew by scaling replichore length with the corresponding skew further improves predictive power. This model outperforms those based solely on genome size, on the longest chromosome, and on the longest replichore (Table 1).

Based on these associations between genome architecture and growth time, we provide some speculative insight on genome evolution.

### Adaptive evolution of bacterial genome architecture

Based on the observed association between genome size and doubling time, we speculate that, within a shared ecological setting, bacteria and archaea with smaller genomes may enjoy a replication-time advantage that confers higher fitness, and such an advantage could favor genome reduction. However, because most bacterial genomes are densely packed with protein-coding genes, any reduction in genome size often entails loss of essential functions, including pathways involved in replication fidelity and DNA repair, as observed in Mycoplasma and other minimal-genome bacteria [23, 24]. Thus, bacterial genomes appear to be shaped by opposing pressures to minimize replication time while maintaining genetic functionality.

Beyond overall size, the correlation with the length of the longest replichore in bacteria highlights the importance of genome organization. Because replication proceeds bidirectionally, the slowest-replicating replichore sets the replication time. Based on the observations, we speculate that bacterial genomes may evolve under pressure not only to reduce size but also to balance replichore lengths, minimizing replication delays and enhancing growth efficiency.

Furthermore, to explore whether the correlation arises from closely related taxa contributing redundant information, we reconstructed ancestral states for the dataset and fitted the model to these reconstructed traits. We found that the model fits the ancestral traits more strongly, with predictive strength (*R*^2^) decreasing progressively along the evolutionary tree as successive speciations occur. Based on this observation, we speculate that with the evolution and increasing complexity of organisms, resulting from genetic drift and other selection pressures, the association between growth rate and genome structure may have been screened.

### Potential adaptive role of compositional skew

Nucleotide compositional skew has traditionally been interpreted either as a neutral footprint of asymmetric mutational processes—such as cytosine deamination on the lagging strand [33, 34]—or as a consequence of selection for gene orientation on the leading strand [34, 35]. Our results suggest an additional possibility: skew may have functional implications for replication dynamics. We find that fast-growing bacteria tend to exhibit stronger purine–pyrimidine asymmetry, and models incorporating skew better predict doubling times. This raises the hypothesis that compositional skew might facilitate faster replication fork progression and thereby contribute to faster growth. Although we observe a modest correlation between skew and experimentally measured replication fork speeds (Fig. 6b), current data are limited, and larger datasets will be required to investigate their relationship.

The influence of skew on replication fork progression speed can be explained using the “Asymmetric cooperative model”, which was originally proposed to explain the evolutionary advantages of unidirectional strand construction during DNA replication and antiparallel orientation of the two strands of dsDNA. An asymmetric kinetic nearest neighbor interaction of base pairs leads to faster strand separation kinetics of a skewed sequence within the above model, thereby facilitating replication and, consequently, becoming evolutionarily advantageous [21, 36]. Wanrooij et al. also showed that simply swapping the two arms of a highly skewed origin sequence, changing the skew profile of the sequence from a ∨-shape to a ∧-shape, abolishes origin functionality [37]. Although this result comes from a non-bacterial system, it demonstrates that changes in compositional asymmetry alone can have mechanistic consequences for replication dynamics. In bacteria, however, any influence of skew on replication fork progression remains speculative and requires direct experimental validation.

### Limitations and future directions

Several simplifications were made in this analysis. The classification of bacteria into free-living versus host-dependent was based on a literature survey and may not fully capture ecological diversity. The assumption that replication fork speed scales linearly with skew is also a crude approximation, as zero skew does not imply zero fork speed. More refined models of fork kinetics, potentially incorporating polymerase dynamics and overlapping rounds of replication under fast growth, are needed. Furthermore, while we used RY cumulative skew to identify replication origins, GC or amino-keto (MK) skew may provide stronger predictions in some genomes. Although the primary outcome, that incorporating skew improves predictive power, was robust across RY, GC, and MK definitions, future work should systematically compare these measures. Understanding the mechanistic basis of these differences will be essential for linking compositional biases to DNA replication dynamics and bacterial fitness.

## Supporting information

TimeEvolutionModel_IV

## Acknowledgments

Support for this work was provided by the Science & Engineering Research Board (SERB), Department of Science and Technology (DST), India, through a Core Research Grant with file no. CRG/2020/003555 and a MATRICS grant with file no. MTR/2022/000086.

## Supplementary Information

The additional analysis performed in this work is provided in Supplementary Information.

The scatter plots of the model-IV fit across 50 time slices are compiled into a video and is provided as TimeEvolutionModel IV.mp4 in the Supplementary Data (GitHub repository, See below).

## Data Availability

The data and the MATLAB script used in this analysis are publicly available at the following GitHub repository:https://github.com/ParthaTbio/Genome_skew_analysis.

## Supplementary Information

This document accompanies the main article “Genome size and nucleotide skews as predictors of bacterial growth rate”. The study investigates the influence of genome size, chromosome organization, replichore length, and nucleotide compositional skew on replication kinetics and, consequently, on bacterial growth rates. Additional analyses on Dataset-I that are not included in the main text are presented here.

### Archaeal doubling time and genome size

The relationship between archaeal genome size (*n* = 153) and doubling time is shown in Supplementary Fig. S1. A moderate correlation (*R*^2^ = 0.12) suggests that genome size contributes to replication kinetics and, consequently, influences growth rate. A stronger correlation is expected when the replichores and their skew are considered rather than the total genome size.

**Supplementary Figure S1:**
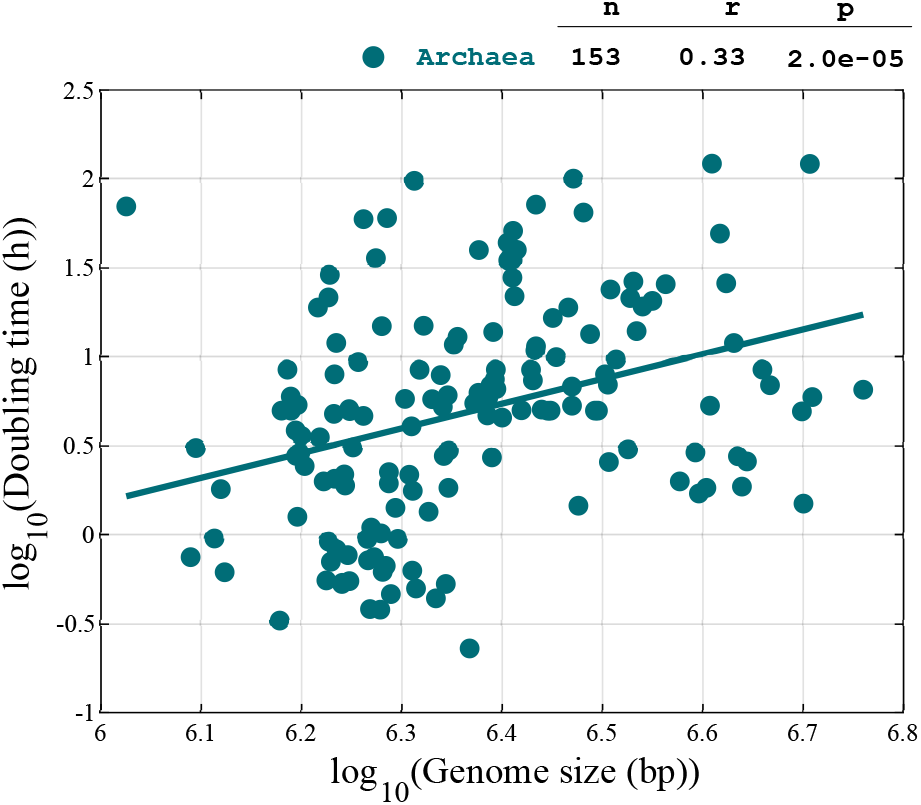
Correlation between archaeal doubling time and total genome size. A moderate association is observed even without accounting for replication parallelization across replichores. A stronger correlation is expected when considering the longest replichore rather than the total genome size.

### Partial correlation analysis

To test whether the observed correlation between the doubling time and the skew normalized replichore length is confounded by other variables, we used the additional data provided in the Dataset-I [13] and performed partial correlation analysis controlling for GC percentage, optimum pH, growth temperature, and optimum temperature, see Supplementary Fig. S2. The analysis is performed on the full data set (Supplementary Fig.S2, left panel) and separately on the free-living ones (Supplementary Fig.S2, right panel). None of these variables shows evidence of confounding the observed association between bacterial doubling time and the skew-normalized replichore length.

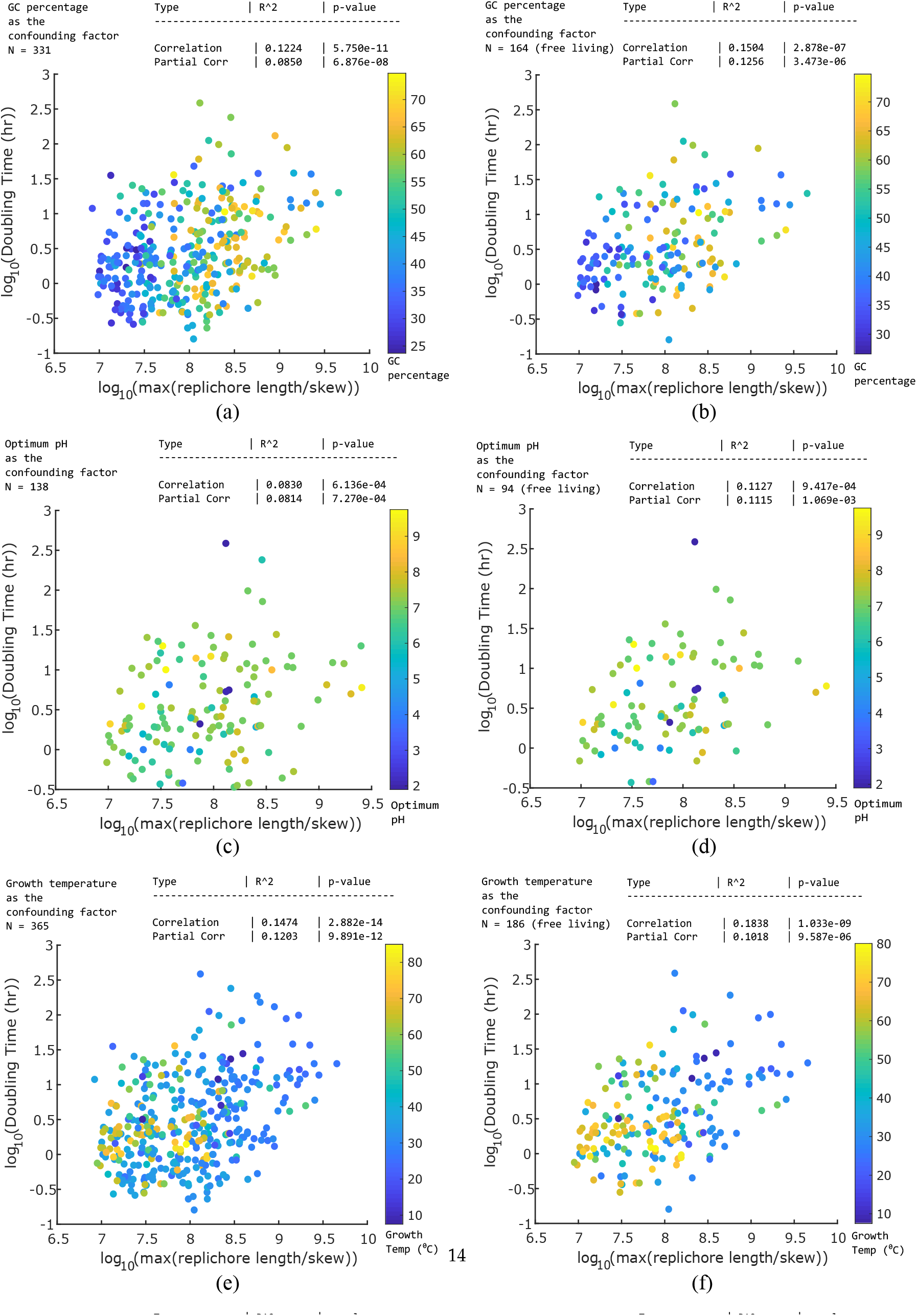

### Intra-clade and inter-clade comparison

Intra-clade correlation was evaluated among species belonging to the same clade, and the results are presented in Supplementary Fig. S4. To compare across clades, we assigned each clade a representative trait value *X*_*j*_ by computing the geometric mean of the trait values of all extant species within that clade. The resulting inter-clade correlations are shown in Supplementary Fig. S3.

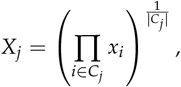

where *X*_*j*_ denotes the trait value assigned to clade *j*, and *C*_*j*_ is the set of species belonging to clade *j*.

**Supplementary Figure S3:**
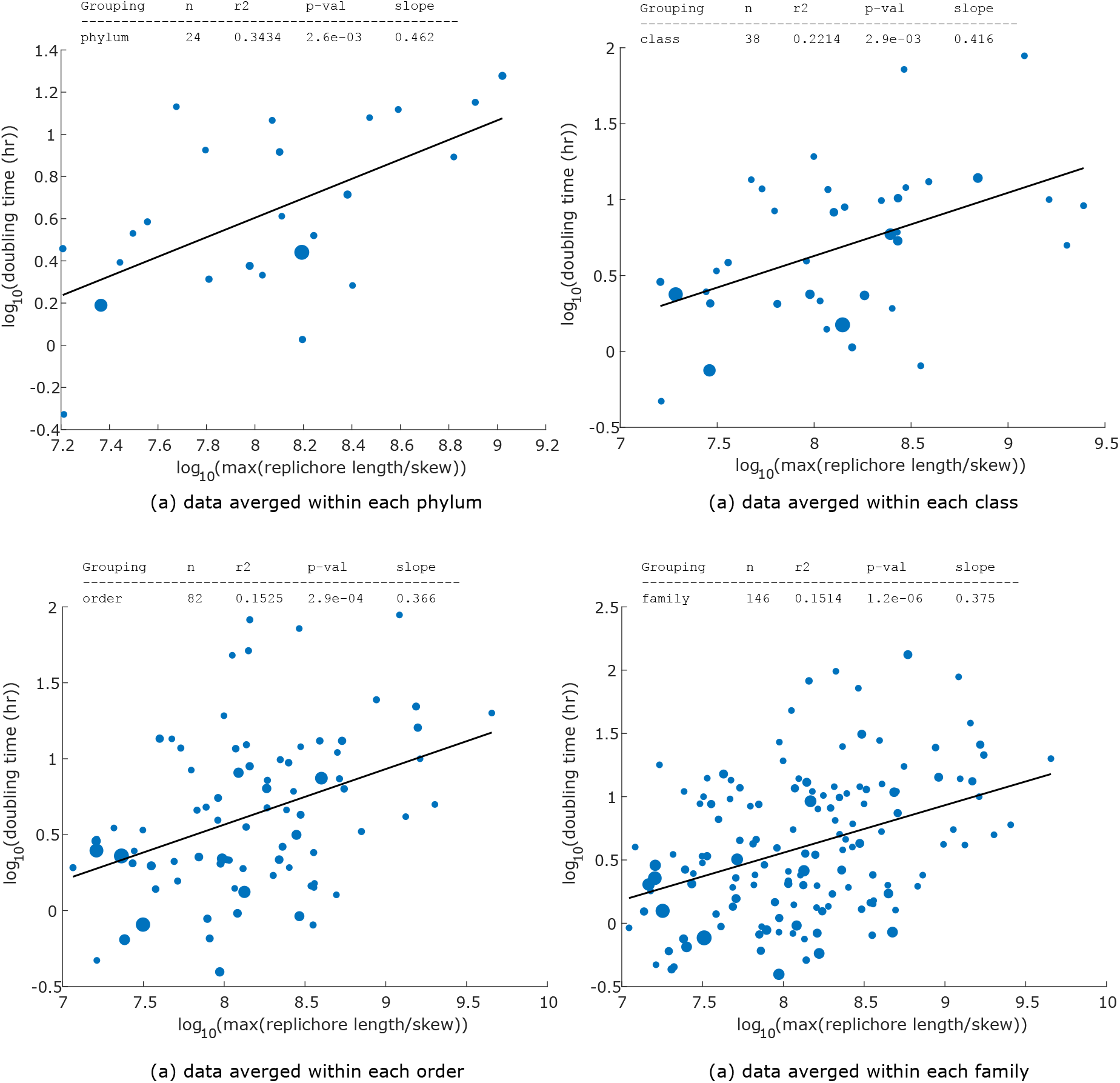
Among clade correlation. The association between skew-normalized replichore length and growth rate becomes progressively stronger, as reflected by increasing *R*^2^, when examined at broader taxonomic levels (family, order, class, phylum). This pattern motivated us to evaluate the model’s fit over deeper evolutionary timescales.

**Supplementary Figure S4:**
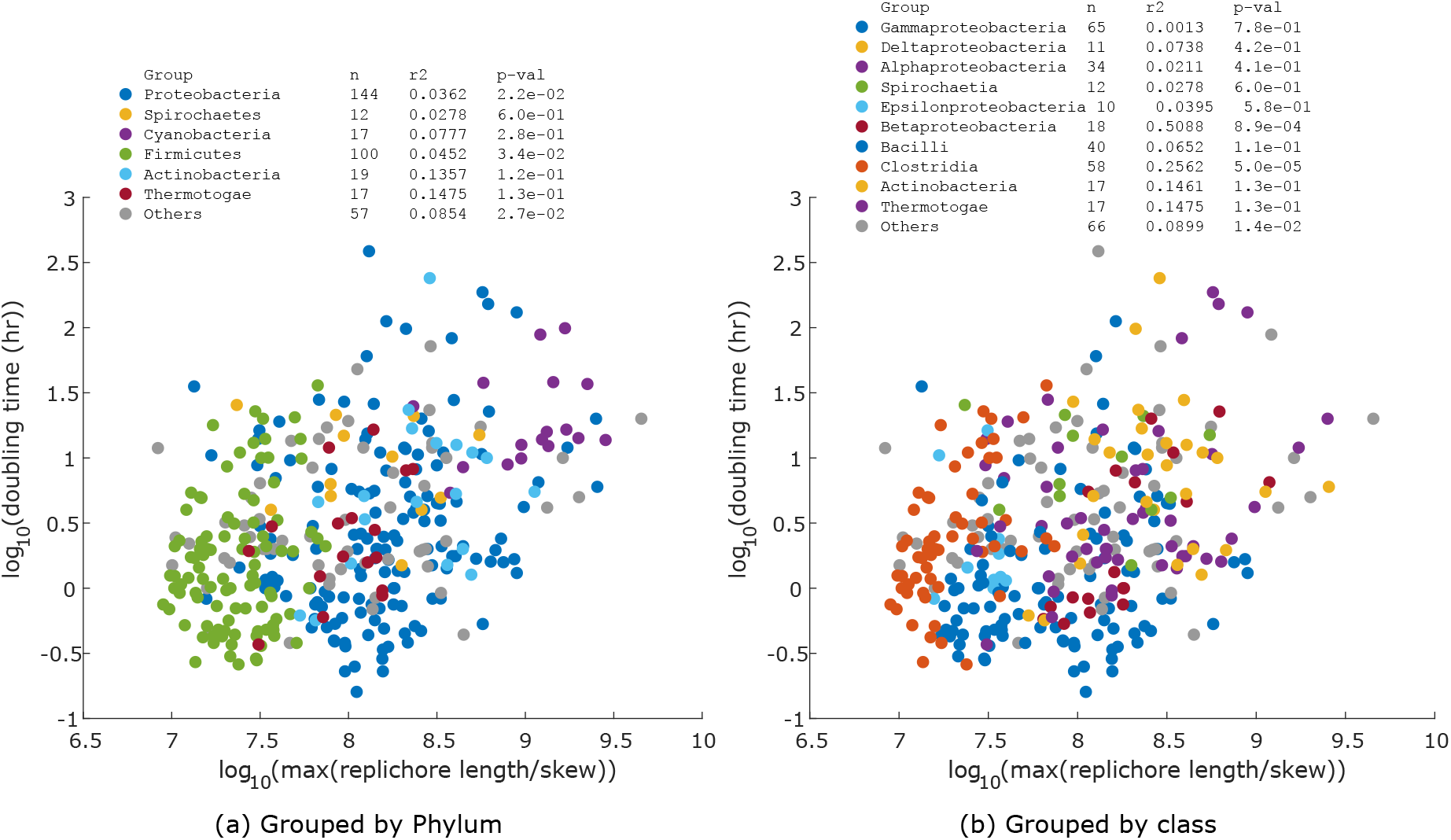
Within clade correlation. The association between skew-normalized replichore length and growth rate is not observed to be stronger within individual clades, with the exception of the classes Clostridia and Betaproteobacteria.

### Stratification Effects on the Correlation

To assess whether the observed correlation could be an artifact of hidden stratification (e.g., a Simpson’s paradox–type effect), we examined the relationship within multiple taxonomic and ecological clusters. We analyzed correlations within Families, Phyla (see Supplementary Fig. S4), metabolic categories, salinity ranges, temperature ranges, and Gram stain groups (see Supplementary Fig. S5). None of these clusters showed an opposite (negative) trend. Thus, the positive association in the full dataset is consistent across diverse subgroups and is unlikely to result from phylogenetic or ecological stratification.

**Supplementary Figure S5:**
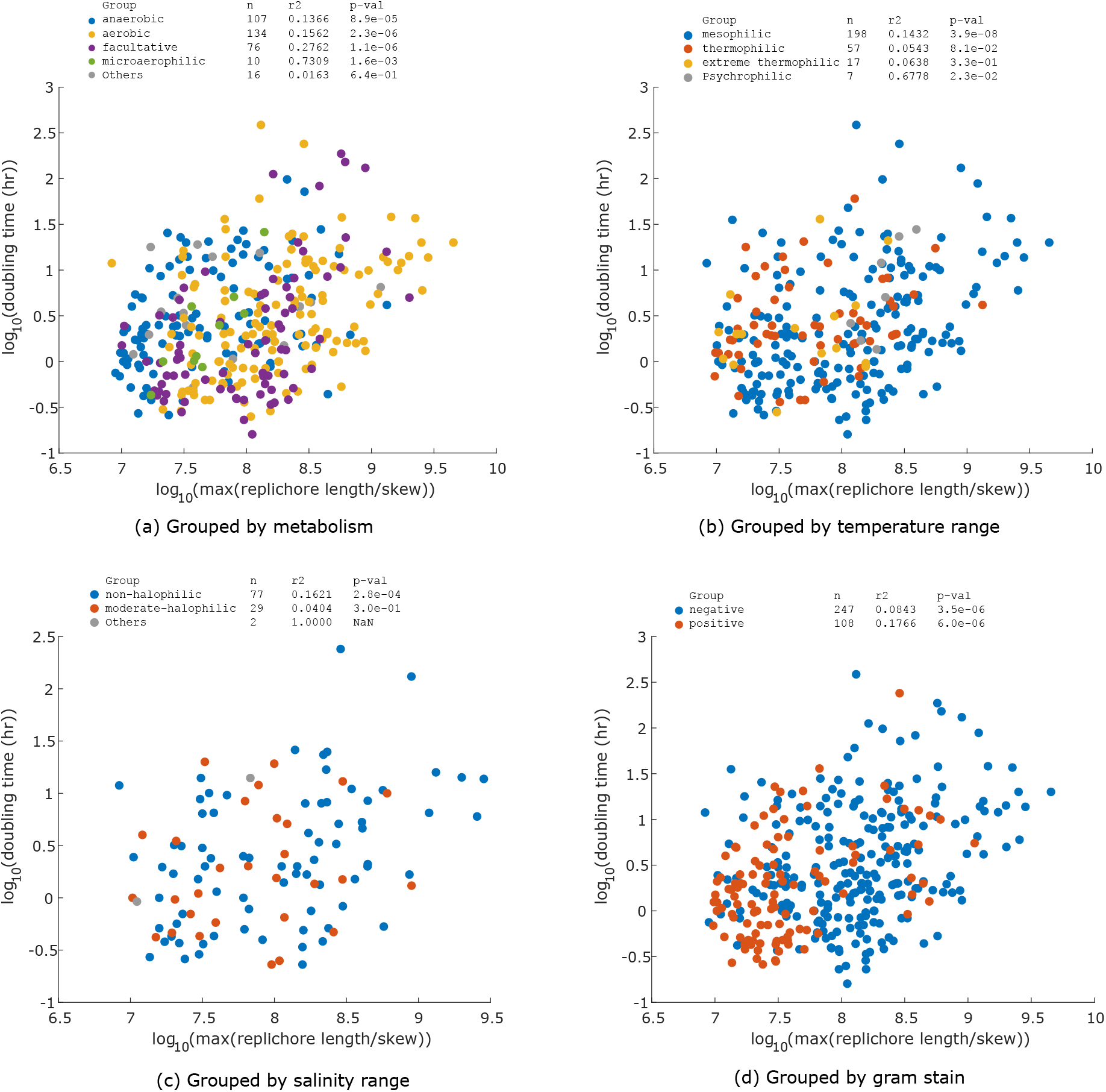
Stratification Effects on the Correlation. Data were clustered by metabolic category, salinity range, temperature range, and Gram stain group. None of these subgroups exhibited an opposite trend to that of the global positive association between skew-normalized replichore length and growth rate. The positive association observed in the full dataset is therefore consistent across diverse ecological and physiological subsets, suggesting that it is unlikely to arise from ecological stratification.

### Phylogenetic tree

The phylogenetic time history data of 323 species of Dataset-I imported from TimeTree [25] is shown in Fig. S6. The tree in Newick format is provided in Supplementary Data.

**Supplementary Figure S6:**
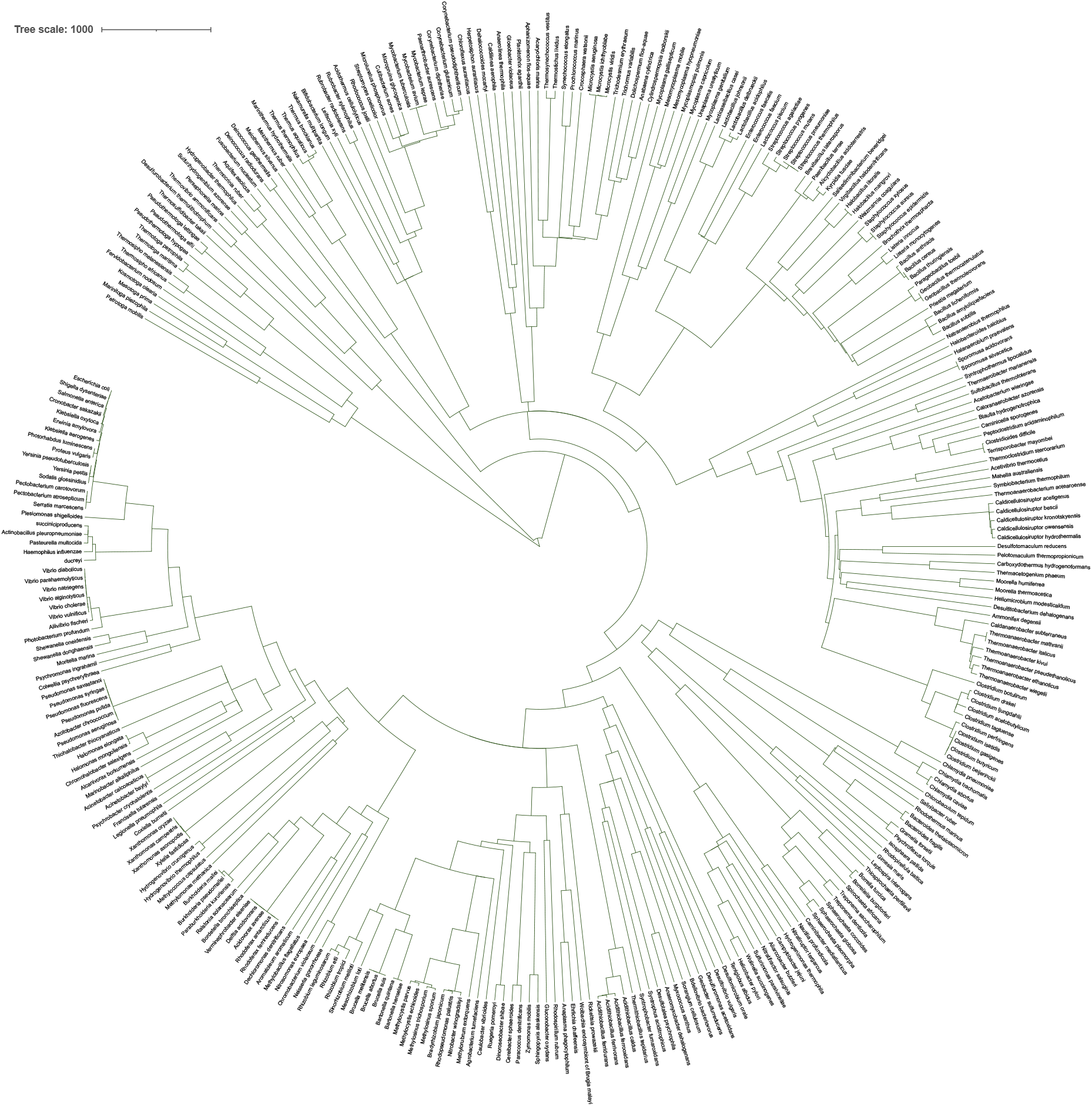
Phylogenetic tree with time history data for 323 species of the Dataset-I imported from TimeTree [25] is shown.

The variance-covariance matrix of the phylogenetic tree is presented as a heat map, Supplementary Fig. S7.

**Supplementary Figure S7:**
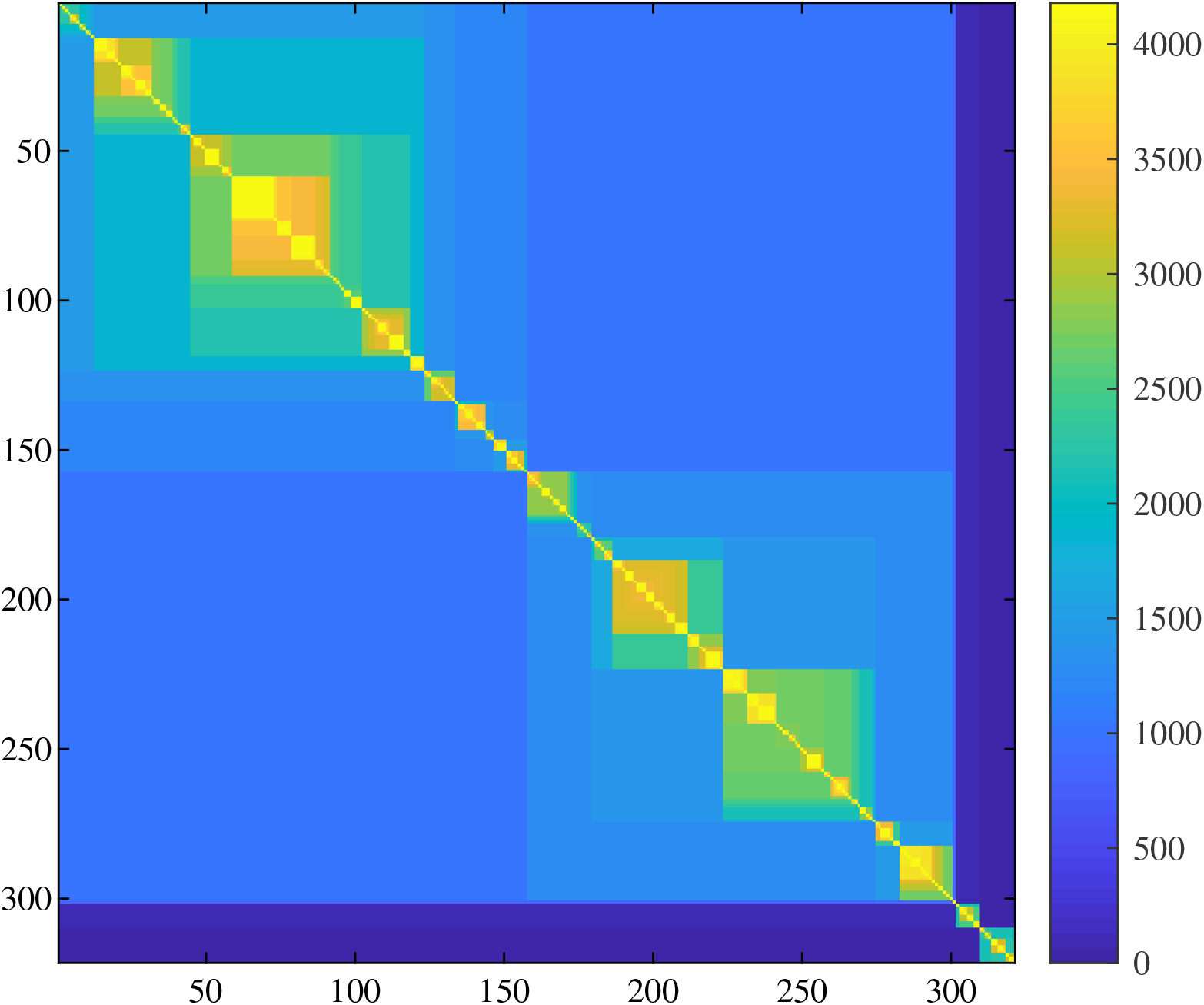
Variance-Covariance matrix corresponding to the phylogenetic tree, after a branch length transformation using Pagel’s *λ* = 0.9.

### Regression Statistics

The regression coefficients *b*_0_ and *b*_1_ representing the slope and intercept of the regression plots, for all 4 models for both Dataset-I and Dataset-II, are provided here in Supplementary Table S1 and Supplementary Table S2, respectively.

**Supplementary Table S1:** Ordinary least squares (OLS) regression statistics for the relationship between doubling time (response) and different genomic predictors (Models I–IV) for Dataset-I. Values shown are coefficient of determination (*R*^2^), sample size (*n*), *p*-value, and regression slope and intercept with their 95% confidence interval (estimate *±* CI).

| Dataset-I | $n$ | $R^2$ | $p$-value | $b_0 \pm \text{SE}$ | $b_1 \pm \text{SE}$ |
| --- | --- | --- | --- | --- | --- |
| **Model I** (Predictor: Genome size) |  |  |  |  |  |
| All | 367 | 0.003 | $2.9 \times 10^{-1}$ | $-0.58 \pm 1.93$ | $0.16 \pm 0.30$ |
| HD+FL | 346 | 0.023 | $4.4 \times 10^{-3}$ | $-2.86 \pm 2.26$ | $0.50 \pm 0.35$ |
| FL only | 187 | 0.095 | $1.8 \times 10^{-5}$ | $-6.10 \pm 2.99$ | $1.02 \pm 0.46$ |
| **Model II** (Predictor: Longest chromosome) |  |  |  |  |  |
| All | 367 | 0.008 | $8.3 \times 10^{-2}$ | $-1.26 \pm 1.95$ | $0.26 \pm 0.30$ |
| HD+FL | 346 | 0.038 | $2.7 \times 10^{-4}$ | $-3.80 \pm 2.26$ | $0.65 \pm 0.35$ |
| FL only | 187 | 0.109 | $3.8 \times 10^{-6}$ | $-6.61 \pm 2.97$ | $1.10 \pm 0.46$ |
| **Model III** (Predictor: Longest replicore) |  |  |  |  |  |
| All | 367 | 0.031 | $7.1 \times 10^{-4}$ | $-2.73 \pm 1.83$ | $0.51 \pm 0.29$ |
| HD+FL | 346 | 0.085 | $3.1 \times 10^{-8}$ | $-5.61 \pm 2.10$ | $0.96 \pm 0.33$ |
| FL only | 187 | 0.152 | $3.7 \times 10^{-8}$ | $-7.42 \pm 2.74$ | $1.27 \pm 0.44$ |
| **Model IV** (Predictor: Skew-normalized replicore) |  |  |  |  |  |
| All | 367 | 0.147 | $2.8 \times 10^{-14}$ | $-2.81 \pm 0.81$ | $0.41 \pm 0.10$ |
| HD+FL | 346 | 0.185 | $5.0 \times 10^{-17}$ | $-3.23 \pm 0.82$ | $0.46 \pm 0.10$ |
| FL only | 187 | 0.183 | $1.1 \times 10^{-9}$ | $-2.80 \pm 1.03$ | $0.42 \pm 0.13$ |

**Supplementary Table S2:** Ordinary least squares (OLS) regression statistics for the relationship between doubling time (response) and different genomic predictors (Models I–IV) for Dataset-II. Values shown are coefficient of determination (*R*^2^), sample size (*n*), *p*-value, and regression slope and intercept with their 95% confidence interval (estimate *±* CI).

| Dataset-II | $n$ | $R^2$ | $p$-value | $b_0 \pm \text{SE}$ | $b_1 \pm \text{SE}$ |
| --- | --- | --- | --- | --- | --- |
| **Model I** (Predictor: Genome size) |  |  |  |  |  |
| All | 180 | 0.040 | $6.9 \times 10^{-3}$ | $3.51 \pm 2.19$ | $-0.47 \pm 0.34$ |
| HD+FL | 159 | 0.008 | $2.5 \times 10^{-1}$ | $-1.10 \pm 2.52$ | $0.23 \pm 0.39$ |
| FL only | 104 | 0.032 | $7.0 \times 10^{-2}$ | $-3.23 \pm 3.94$ | $0.55 \pm 0.60$ |
| **Model II** (Predictor: Longest chromosome) |  |  |  |  |  |
| All | 180 | 0.028 | $2.6 \times 10^{-2}$ | $3.04 \pm 2.25$ | $-0.40 \pm 0.35$ |
| HD+FL | 159 | 0.018 | $8.8 \times 10^{-2}$ | $-1.85 \pm 2.55$ | $0.34 \pm 0.39$ |
| FL only | 104 | 0.062 | $1.1 \times 10^{-2}$ | $-4.73 \pm 3.92$ | $0.78 \pm 0.60$ |
| **Model III** (Predictor: Longest replicore) |  |  |  |  |  |
| All | 180 | 0.010 | $1.8 \times 10^{-1}$ | $1.98 \pm 2.18$ | $-0.24 \pm 0.35$ |
| HD+FL | 159 | 0.045 | $7.1 \times 10^{-3}$ | $-2.94 \pm 2.39$ | $0.53 \pm 0.38$ |
| FL only | 104 | 0.113 | $4.8 \times 10^{-4}$ | $-6.02 \pm 3.53$ | $1.02 \pm 0.56$ |
| **Model IV** (Predictor: Skew-normalized replicore) |  |  |  |  |  |
| All | 180 | 0.042 | $5.6 \times 10^{-3}$ | $-1.36 \pm 1.30$ | $0.23 \pm 0.16$ |
| HD+FL | 159 | 0.204 | $2.3 \times 10^{-9}$ | $-3.38 \pm 1.17$ | $0.46 \pm 0.14$ |
| FL only | 104 | 0.300 | $1.8 \times 10^{-9}$ | $-4.43 \pm 1.46$ | $0.59 \pm 0.18$ |

### Testing for potential outliers

Comparison of ordinary least squares (OLS) regression and OLS refined statistics on Dataset-I for the relationship between skew-normalized replichore length (predictor) and doubling time (response) is shown in Supplementary Table S3. “OLS-refined” denotes OLS refitted after excluding influential data points with Cook’s distance *D*_*i*_ > 4/(*n* − *k* − 1), where *n* is the sample size and *k* is the number of independent variables (*k* = 1 in this analysis). The consistency in *R*^2^ and *p*-value suggests the robustness of the OLS, indicating that the regression is not biased by a single or a few data points.

**Supplementary Table S3:**
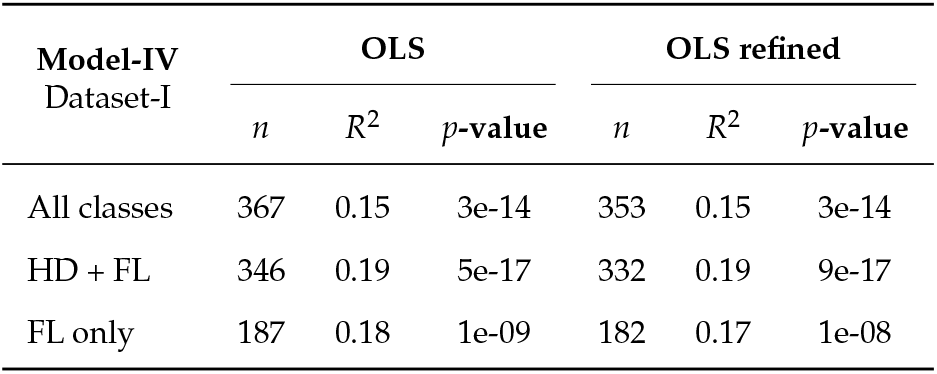
Comparison of ordinary least squares (OLS) regression and OLS-refined statistics on Dataset-I for the relationship between skew-normalized replichore length (predictor) and doubling time (response).

| Model-IV<br>Dataset-I | OLS |  |  | OLS refined |  |  |
| --- | --- | --- | --- | --- | --- | --- |
| | $n$ | $R^2$ | $p$-value | $n$ | $R^2$ | $p$-value |
| All classes | 367 | 0.15 | 3e-14 | 353 | 0.15 | 3e-14 |
| HD + FL | 346 | 0.19 | 5e-17 | 332 | 0.19 | 9e-17 |
| FL only | 187 | 0.18 | 1e-09 | 182 | 0.17 | 1e-08 |

